# Exploration of the *Ixodes ricinus* virosphere unveils an extensive virus diversity including novel Coltiviruses and other reoviruses

**DOI:** 10.1101/2021.04.08.438920

**Authors:** Bert Vanmechelen, Michelle Merino, Valentijn Vergote, Lies Laenen, Marijn Thijssen, Joan Martí-Carreras, Edwin Claerebout, Piet Maes

## Abstract

Recent metagenomics studies have revealed several tick species to host a variety of previously undiscovered RNA viruses. *Ixodes ricinus*, which is known to be a vector for many viral, bacterial and protozoan pathogens, is the most prevalent tick species in Europe. For this study, we decided to investigate the virosphere of Belgian *I. ricinus* ticks. High-throughput sequencing of tick pools collected from six different sampling sites revealed the presence of viruses belonging to many different viral orders and families, including *Mononegavirales, Bunyavirales, Partitiviridae* and *Reoviridae*. Of particular interest was the detection of several new reoviruses, two of which cluster together with members of the genus *Coltivirus*. This includes a new strain of Eyach virus, a known causative agent of tick-borne encephalitis. All genome segments of this new strain are highly similar to those of previously published Eyach virus genomes, except for the fourth segment, encoding VP4, which is markedly more dissimilar, potentially indicating the occurrence of a genetic reassortment. Further PCR-based screening of over 230 tick pools for 14 selected viruses showed that most viruses could be found in all six sampling sites, indicating the wide spread of these viruses throughout the Belgian tick population. Taken together, these results illustrate the role of ticks as important virus reservoirs, highlighting the need for adequate tick control measures.

## Introduction

Recent metagenomics studies have revealed the extensive virus diversity within arthropods [1,2]. Of particular interest amongst arthropods are the hematophagous vectors. These animals feed on the blood of vertebrate hosts, presenting the ideal conditions for the transmission of pathogens. In addition to mosquitoes, ticks are some of the most important vectors of infectious diseases, both in humans, livestock and wildlife animals [3]. Ticks are known to carry a diverse range of pathogens, including protozoa, bacteria, helminths and viruses, although in Europe they are predominantly known for their ability to transmit *Borrelia burgdorferi* bacteria, the causative agent for Lyme disease [4-6]. However, ticks also act as vectors for multiple viral pathogens, including tick-borne encephalitis virus (TBEV), Crimean-Congo haemorrhagic fever virus and Colorado tick fever virus [7]. In the last decade, many articles have been published looking at the virus diversity in different tick species in different regions, revealing the presence of many previously undiscovered viruses [8-14]. Although estimating the relative risk of these novel viruses for human, animal and plant health is difficult, their phylogenetic clustering within specific virus families and genera that are known to harbour tick-borne human, animal and plant pathogens (*Reoviridae, Nairoviridae, Flaviviridae, Phenuiviridae* …) hints at pathogenic potential for at least some of them.

Ticks are divided into three families: the *Argasidae* or ‘soft ticks’, the *Ixodidae* or ‘hard ticks’ and the *Nuttalliellidae*. Unlike the latter, which contains only one species, the families *Argasidae* and *Ixodidae* harbour several hundred member species, split across multiple genera [15]. In Belgium, only a limited number of tick species has been described and almost all reported tick bites can be attributed to the castor bean tick or *Ixodes ricinus*, the type species of the genus *Ixodes* in the family of ‘hard ticks’ [16,17]. This predominantly European tick species requires three blood meals during its life cycle: to moult from larva to nymph and from nymph to adult, and to lay eggs once it has reached adulthood [18]. This triple-host life cycle makes *I. ricinus* an important vector for the transmission of several tick-borne diseases between a variety of mammalian hosts, including humans. Several viruses that pose a threat to human and animal health are already known to be transmitted by *I. ricinus* ticks, although, with the exception of TBEV, studies concerning human diseases of viral origin spread by *I. ricinus* are limited (see [19] for a detailed overview). Furthermore, the ongoing discovery of new (putative) viral pathogens, such as the newly described Alongshan virus, which was recently reported to be present in European *I. ricinus* ticks, indicates that *I. ricinus* ticks might harbour more pathogenic viruses than hitherto assumed [20].

In recent years, a number of studies have delved deeper into the virus diversity in *I. ricinus* in Europe, revealing the presence of several previously undetected viruses [10,20-22]. In 2017, we reported the discovery of a novel nairovirus, Grotenhout virus, in Belgian *I. ricinus*, which was identified through next-generation sequencing of a pool of adult ticks [23]. Because we also detected traces of other viruses, we decided to sequence additional tick pools to further characterize the virus diversity in Belgian ticks. Here we report the detection of several novel as well as previously described viruses, including a rhabdovirus and multiple reoviruses. Additionally, we performed PCR screening for 14 different viruses, including the novel viruses reported here as well as some other viruses known to be carried by *I. ricinus*, on more than 200 tick pools from six sampling locations in Belgium, providing a broad overview of virus prevalence in Belgian *I. ricinus* ticks.

## Materials and methods

### Sample collection

Ticks were collected between 2009-2017 from six sampling locations in Belgium (Torhout, Moerbeke, Zoersel, Gierle, Chimay and Heverlee), by flagging the low-growing vegetation. Ticks were kept at - 20°C or in 100% ethanol for long-term storage. A total of 767 ticks were collected, including 33 adults, 513 nymphs and 221 larvae. All ticks were sorted based on their location of origin and their developmental stage and all were identified as *I. ricinus* based on phenotypical characteristics. An overview of the different sampling locations and the number of ticks caught at each location can be found in Figure 1.

**Figure 1:**
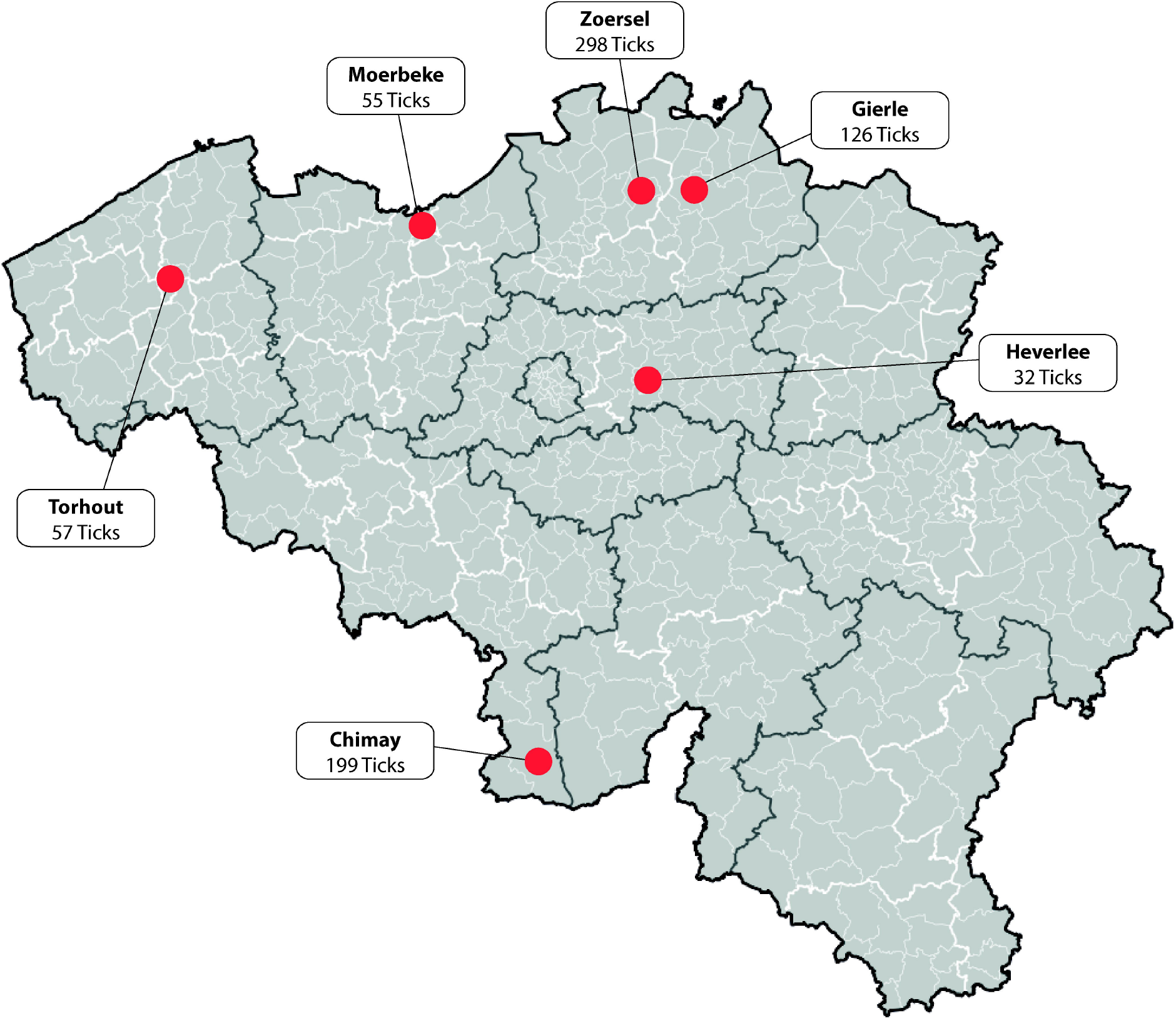
Tick sampling sites. Ticks were collected from six sampling sites in Belgium, marked in red.

### RNA extraction

Each adult tick or pool (1-10) of nymphs/larvae (see Supplementary Table S1) was added to a tube containing 100 µl PBS and zirconium oxide beads and subsequently homogenized for 150s at 4000g in a Bertin Minilys homogenizer. Total RNA was extracted from the homogenate using the RNeasy Mini Kit (QIAGEN, Leiden, Netherlands) according to the manufacturer’s instructions. DNA carry-over was further depleted by DNase treatment using 2U TurboDNase (Invitrogen, ThermoFisher Scientific, California, USA) and the remaining RNA was re-purified using the RNeasy MinElute Cleanup Kit (QIAGEN, Leiden, Netherlands).

### Nanopore sequencing

RNA extract from a single female adult tick was converted to cDNA using the Complete Whole Transcriptome Amplification Kit (WTA2, Sigma-Aldrich, Saint Louis, MO, USA). Purification of cDNA was done using the MSB Spin PCRapace kit (Stratec Molecular, Berlin, Germany). The resulting cDNA was prepared for nanopore sequencing using the SQK-LSK108 kit (Oxford Nanopore Technologies, Oxford, UK) and loaded on an R9.4.1 flow cell. Reads corresponding to tick rRNA were removed and, following size selection, reads between 200-2,000 nucleotides were assembled using Canu v1.5 [24]. Identification of the resulting contigs was done using BLAST (https://blast.ncbi.nlm.nih.gov/Blast.cgi).

### Illumina sequencing

Ten pools were made for Illumina sequencing, containing 3-5 RNA extracts each. An overview of the ten pools can be found in Table 1. RNA was converted to cDNA using the WTA2 Kit (Sigma-Aldrich). cDNA was purified using the MSB Spin PCRapace kit (Stratec Molecular). Sequencing libraries were prepared using the Illumina NexteraXT DNA Library Prep Kit (Illumina, San Diego, USA). Libraries were double indexed using Set A-barcoded adapters (Illumina, San Diego, USA). In-house modifications on the standard library preparation were conducted following Conceicão-Neto et al. [25]. Sequencing was performed on a HiSeq4000 platform (Illumina, San Diego, USA) at VIB – Nucleomics Core (Leuven, Belgium). CLC Genomics Workbench (v10.1.1) was used to trim the reads by adapter content, quality (Phred q-score >35) and length (>25 bp), and to assemble the resulting trimmed reads. Diamond (v0.9.24) was used to classify the obtained contigs, using the blastx algorithm against the complete NCBI non-redundant protein database [26].

**Table 1:** Illumina sequencing data overview.

| Pool number | RNA pools used* |  | #Ticks | Adult/Nymph/Larva | Location | #Reads | #Viral reads | #Contigs | #Viral contigs |
| --- | --- | --- | --- | --- | --- | --- | --- | --- | --- |
| S1 | 142<br>144 | 147<br>150 | 4 | 4 Adults | Moerbeke | 62,886,446 | 977 | 4,874 | 13 |
| S2 | 4<br>6 | 87<br>91 | 4 | 4 Adults | Chimay<br>Gierle | 65,898,424 | 1,576 | 6,918 | 16 |
| S3 | 140<br>162 | 168 | 3 | 3 Adults | Heverlee<br>Torhout | 48,976,538 | 6,621 | 25,558 | 24 |
| S4 | 143<br>146 | 149 | 3 | 3 Adults | Moerbeke | 68,520,456 | 12,952 | 65,273 | 65 |
| S5 | 89<br>90 | 95 | 3 | 3 Adults | Gierle | 68,966,000 | 750,760 | 6,771 | 94 |
| S6 | 129<br>159 | 161 | 7 | 3 Nymphs<br>4 Larvae | Heverlee<br>Moerbeke | 63,439,638 | 149,654 | 74,219 | 693 |
| S7 | 47<br>48 | 51<br>56 | 12 | 12 Nymphs | Chimay | 76,725,768 | 7,301 | 15,224 | 66 |
| S8 | 107<br>110 | 114<br>126 | 12 | 12 Nymphs | Chimay<br>Gierle | 58,129,700 | 366,933 | 53,572 | 132 |
| S9 | 106<br>110<br>111 | 112<br>113 | 15 | 15 Nymphs | Gierle | 83,201,118 | 293,361 | 35,840 | 842 |
| S10 | 39<br>44<br>45 | 135<br>139 | 13 | 13 Nymphs | Chimay<br>Heverlee | 70,097,012 | 47,741 | 15,718 | 96 |

\*See Supplementary Table S1

### PCR screening and genome completion

The Qiagen OneStep RT-PCR kit was used to screen for the different viruses, and to complete gaps in the obtained genome sequences. The following cycling conditions were used: 50°C for 30 min, 95°C for 15 min, 40 cycles of 94°C for 30 s, a primer-specific annealing temperature for 30 s and 72°C for 60 s, and a final extension step at 72°C for 10 min. Completion of genome ends was done using the Roche 5’/3’ RACE kit, 2^nd^ generation. An overview of all used screening primer sets and annealing temperatures can be found in Supplementary Table S2. PCR products were visualized by 2% agarose gel electrophoresis and Sanger sequenced to confirm their identity. Sequencing was done by Macrogen Europe (Macrogen Europe B.V., Amsterdam, The Netherlands) after the products were purified in-house with PureIT ExoZAP (Ampliqon, Odense, Denmark).

### Phylogenetic analysis

Multiple sequence alignments were made based on deduced amino acid sequences of the RNA-dependent RNA polymerase (RdRp) sequence of members of the families *Rhabdoviridae* and *Reoviridae*. For the rhabdovirus phylogenetic tree, amino acid sequences for all currently recognized International Committee on the Taxonomy of Viruses (ICTV) species, supplemented with all unclassified rhabdoviruses >5,500 bp (NCBI: txid35303) were downloaded from NCBI-Nucleotide (https://www.ncbi.nlm.nih.gov/nuccore). Seqkit was used to keep only protein sequences >1,800 amino acids, which were aligned using MAFFT (v7.310; localpair algorithm) [27,28]. Based on this alignment, the closest related sequences to Chimay rhabdovirus (and Rabies virus as an outgroup) were realigned using MAFFT. The resulting alignment was trimmed using TrimAl (v1.4.rev15; gappyout setting), and model selection and phylogenetic tree calculation were done using IQ-TREE (v.1.6.12), employing 1000 bootstrap replicates [29,30]. The resulting tree was visualized using Figtree v1.4.3. For the reovirus tree, the RdRp protein sequence was deduced from all (putative) members of the subfamily *Spinareovirinae* for whom an RdRp-encoding segment could be identified. Alignment generation, trimming and phylogenetic tree calculation and visualization were performed as described above. An overview of the accession numbers of all used sequences can be found in Supplementary Table S4.

## Results

### Detection of Chimay rhabdovirus

We reported previously the discovery of a novel nairovirus, Grotenhout virus, in a pooled library of 10 adult tick RNA extracts [23]. Even though we recovered a complete L and S segment from the Illumina data, we found no traces of an M segment. However, we did find traces of a novel phlebovirus, which we provisionally named Leuven phlebovirus (GenBank ID: MG407659-MG407660). In an attempt to obtain more sequence information for this novel phlebovirus and to simultaneously look for the missing Grotenhout virus M segment, we screened all ten RNA extracts used to make the Illumina pool for both viruses by RT-PCR, and selected one dual-positive extract (pool 7; see Supplementary Table S1) for RNA sequencing using the Oxford Nanopore MinION. This run yielded ∼4.6 million reads, covering ∼1.81 Gb. Following rRNA removal and length trimming (keeping reads between 200-2,000 bp), assembly using Canu (v1.5) yielded 31 contigs. Together, these contigs cover the full-length of both the Grotenhout virus L and S segments, as well as the L and S segments of the phlebovirus. All four segments had excellent coverage (see Table 2), but for neither virus a contig matching a putative M segment was detected. The remaining contigs could all be classified as being of host or bacterial origin, except for three contigs that displayed limited similarity with members of the family *Rhabdoviridae* and appeared to be part of the same, novel rhabdovirus. Correction by Sanger sequencing and RACE was used to obtain the complete 13,706 bp genome sequence for this virus, which was given the name Chimay rhabdovirus (GenBank ID: MF975531). The genome has the typical rhabdovirus genome architecture, encoding five open reading frames (ORF): 3’-N-P-M-G-L-5’ and clusters alongside Blacklegged tick rhabdovirus 1 (Figure 2).

**Table 2:**
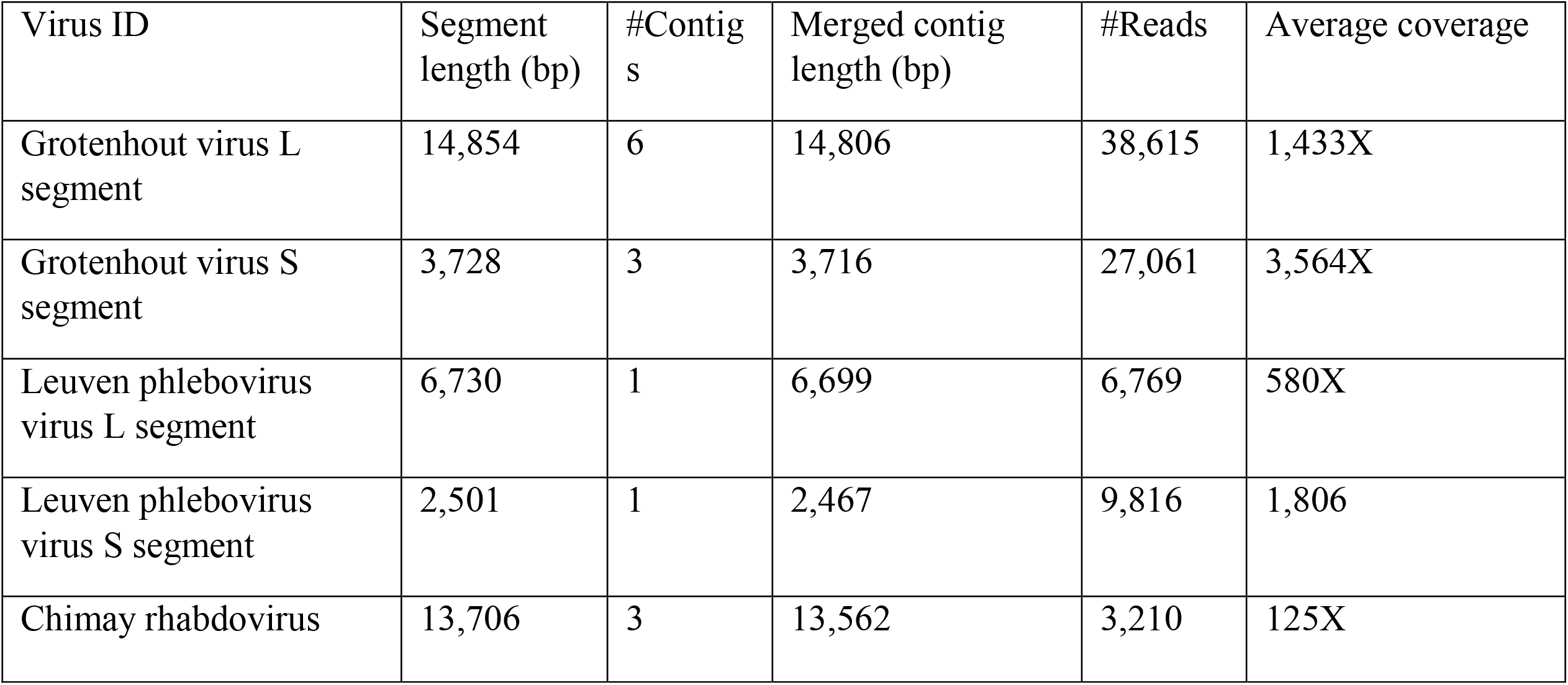
Tick pool 7 nanopore data.

| Virus ID | Segment length (bp) | #Contigs | Merged contig length (bp) | #Reads | Average coverage |
| --- | --- | --- | --- | --- | --- |
| Grotenhout virus L segment | 14,854 | 6 | 14,806 | 38,615 | 1,433X |
| Grotenhout virus S segment | 3,728 | 3 | 3,716 | 27,061 | 3,564X |
| Leuven phlebovirus virus L segment | 6,730 | 1 | 6,699 | 6,769 | 580X |
| Leuven phlebovirus virus S segment | 2,501 | 1 | 2,467 | 9,816 | 1,806 |
| Chimay rhabdovirus | 13,706 | 3 | 13,562 | 3,210 | 125X |

**Figure 2:**
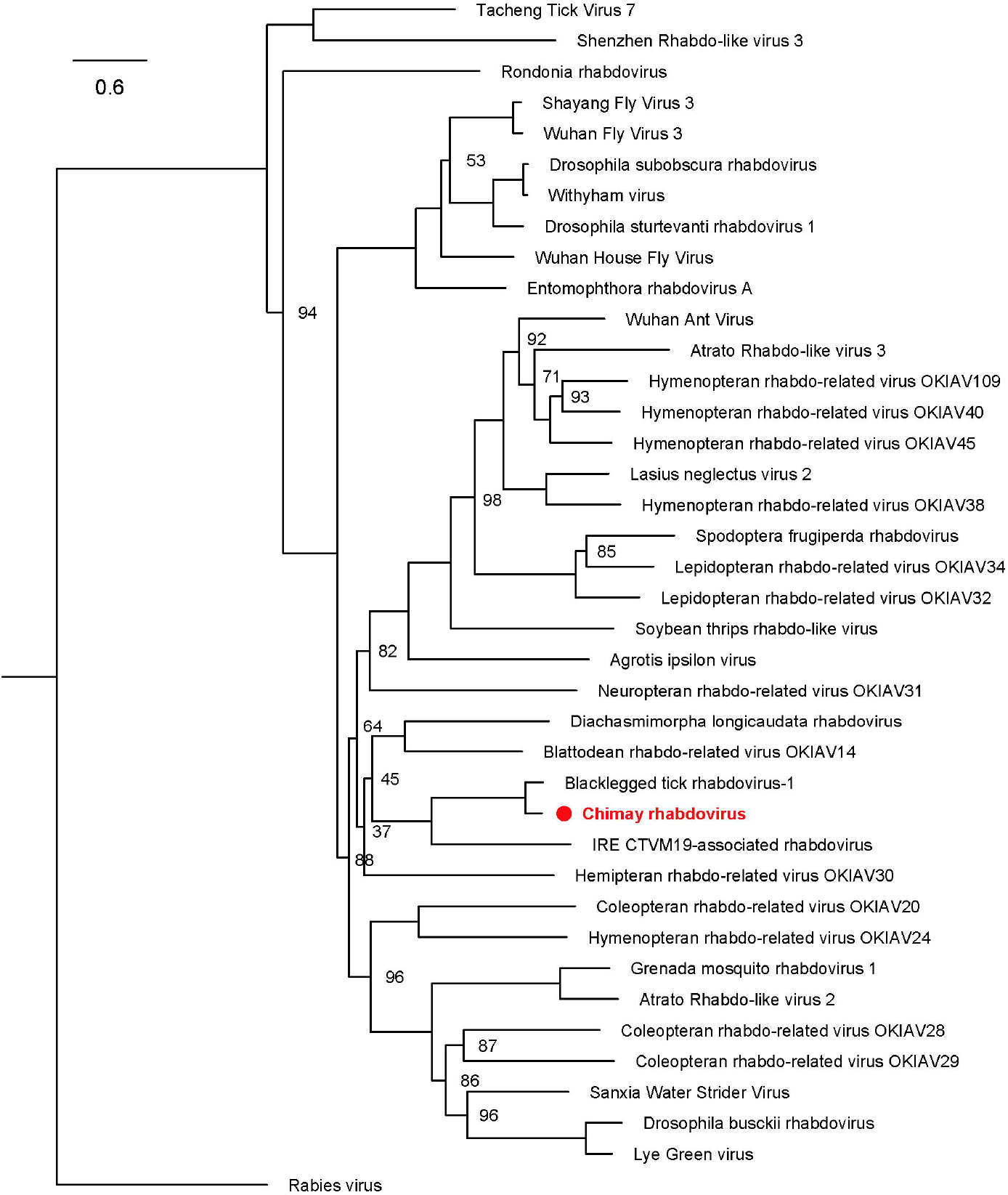
Phylogenetic clustering of Chimay rhabdovirus. Maximum-likelihood tree based on the protein sequence of the RdRp of Chimay rhabdovirus and the closest-related (putative) members of the family *Rhabdoviridae*, including Rabies virus as an outgroup. Chimay rhabdovirus clusters near Blacklegged tick rhabdovirus-1 and IRE CTVM19-associated rhabdovirus, both tick-derived viruses. Branch lengths are drawn according to scale, representing the amount of amino acid substitutions. The numbers at the node indicate the bootstrap support for each node, based on 1000 replicates. Only values <100 are shown. All used accession numbers are provided in Supplementary Table S4.

### Tick sample set

The finding of three separate viruses in the RNA isolated from a single tick indicated that ticks harbour more viruses than previously assumed. To further investigate this, we decided to collect additional ticks of different developmental stages and from different locations in Belgium and used next-generation sequencing for the detection of known and novel viruses in these ticks. Ten tick pools, made from pooling 3–5 RNA extracts, were sequenced on an Illumina 4000 HiSeq, yielding 48,976,538– 83,201,118 reads per pool. Following adapter trimming, between 4,874–74,219 contigs could be assembled for each pool (Table 1). Although the fraction of viral reads compared to host material was low for most pools, we did detect traces of many different viruses, with findings differing strongly from pool to pool (Figure 3). While some pools were largely dominated by the presence of one specific virus (Pool 5), others displayed strong diversity, containing traces of viruses belonging to many different families. Interestingly, while most of the virus families identified in our data are known to contain tick-borne viruses, we also observed fungus/plant-specific viral families, especially in Pool 6. This pool also displayed the largest diversity, but this is most likely attributable to the presence of fungi in the sample used for RNA extraction. Whether these fungi were infecting the ticks in this sample or should be considered environmental contaminants remains to be determined.

**Figure 3:**
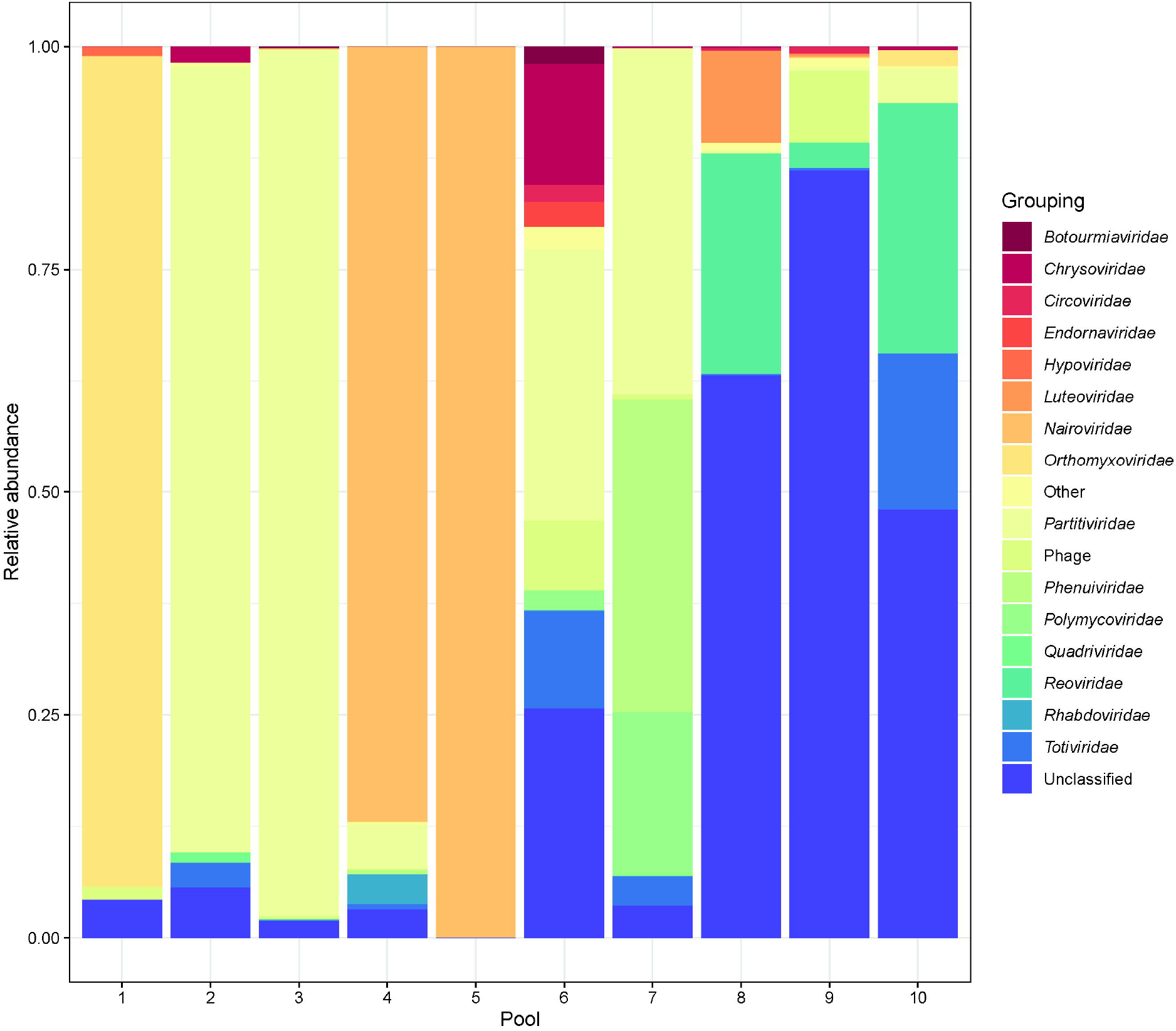
Abundance of viral families in the different Illumina pools. Fraction of viral reads per pool, color-coded based on the taxonomical classification of the closest available reference, as determined by DIAMOND. Taxonomical groups representing <1% of the virus reads in a sample are grouped as ‘Other’. Virus families known to be families of bacteriophages are grouped together as ‘Phage’, as they are unlikely to represent true tick-associated viruses.

### Reconstruction of viral genomes

While for most detected viruses only fragments of the genome were retrieved, some were present more abundantly, allowing the reconstruction of near-complete genome sequences. These include Grotenhout virus and the aforementioned Leuven phlebovirus. Similar isolates have recently also been discovered in northern European ticks, under the name Norway phlebovirus 1 [11]. Several other viruses that were discovered in this study are also abundantly present in our data, including Bronnoya virus, Norway partiti-like virus 1 and Norway luteo-like virus 1. Lastly, we also managed to retrieve near-complete genome sequences of three reoviruses. These include a novel strain of Eyach virus, named strain Heverlee (GenBank ID: MW874081-MW874092), as well as two previously undescribed viruses, named Gierle tick virus (GenBank ID: MW874093-MW874104) and Zoersel tick virus (GenBank ID: MW874105-MW874114), respectively. For Eyach virus strain Heverlee, a combination of Illumina and Sanger sequencing was used to obtain the complete sequence of all 12 genome segments. While almost all segments are highly similar (>94% nucleotide identity) to previously identified strains of Eyach virus (Table 3), the fourth segment, encoding VP4, is markedly dissimilar, displaying <78% nucleotide similarity. Akin to the Eyach virus, we managed to identify all 12 genome segments of Gierle tick virus, a putative new viral species in the genus *Coltivirus* most similar to Tarumizu tick virus and Kundal virus (Figure 4). Lastly, we also managed to retrieve the near-complete genome of a third reovirus, Zoersel tick virus. Although markedly dissimilar from all currently classified reoviruses, this virus shares some similarity with the unclassified reoviruses High Island virus, Eccles virus and ‘Reoviridae sp. BF02/7/10‘, and the unclassified Riboviria Shelly beach virus and Hubei diptera virus 21. Together, these five viruses form a separate clade within the family *Reoviridae*, and for all five, only six genome segments are known, setting them apart from all other reoviruses, which typically have 10-12 genome segments. For Zoersel tick virus, we initially also identified only six genome segments using BLAST-based approaches. However, by reducing the stringency of our significance thresholds and by verifying putative hits with profile hidden Markov models using HMMER v3.3.2, an additional three segments could be identified that shared similarity with Operophtera brumata reovirus, a 10-segmented, unclassified reovirus that phylogenetically clusters as a sister-clade to the aforementioned 6-segmented viruses [31,32]. By looking through the unannotated contigs of our Illumina data, a putative tenth segment was identified for Zoersel tick virus. This contig shows no significant similarity to any known sequence (viral or otherwise), but was present in both Illumina pools containing Zoersel tick virus (and only these pools) and was the only unannotated contig in both sequencing pools that had comparable read coverage to the contigs of the nine segments that could be annotated (Supplementary Table S3). Its length also corresponds to the presumed length of the missing segment and the sequence is predicted to encode one large ORF, as is characteristic for reovirus segments. Therefore, Zoersel tick virus is believed to have 10 genome segments, akin to the distantly related Operophtera brumata reovirus. In addition to these three reoviruses for which (near-)complete genome could be retrieved, we also detected traces of a fourth reovirus in one of the pools: Torhout tick virus (GenBank ID: MW874115-MW874117), a colti-like virus distantly related to the recently discovered Fennes virus, but only partial sequences could be reconstructed for this virus.

**Table 3:** Comparison of Eyach virus (RefSeq) and Eyach virus strain Heverlee.

| Segment | Encoded protein | Segment length |  | Nucleotide identity (%) | Amino acid identity (%) |
| --- | --- | --- | --- | --- | --- |
|  |  | Eyach virus | Heverlee tick virus |  |  |
| 1 | VP1 (RdRp) | 4349 | 4349 | 99.13 | 99.30 |
| 2 | VP2 | 3934 | 3934 | 95.02 | 98.51 |
| 3 | VP3 | 3585 | 3585 | 99.25 | 99.66 |
| 4 | VP4 | 3156 | 3156 | **77.96** | **88.12** |
| 5 | VP5 | 2398 | 2398 | 94.75 | 97.87 |
| 6 | VP6 | 2178 | 2178 | 94.44 | 96.14 |
| 7 | VP7 | 2139 | 2139 | 95.65 | 96.70 |
| 8 | VP8 | 2028 | 2028 | 97.49 | 98.48 |
| 9 | VP9' | 1884 | 1884 | 97.19 | 99.17 |
|  | VP9 |  |  |  | 99.41 |
| 10 | VP10 | 1879 | 1879 | 94.89 | 97.02 |
| 11 | VP11 | 1002 | 1002 | 99.40 | 99.35 |
| 12 | VP12 | 678 | 678 | 95.87 | 94.57 |

**Figure 4:**
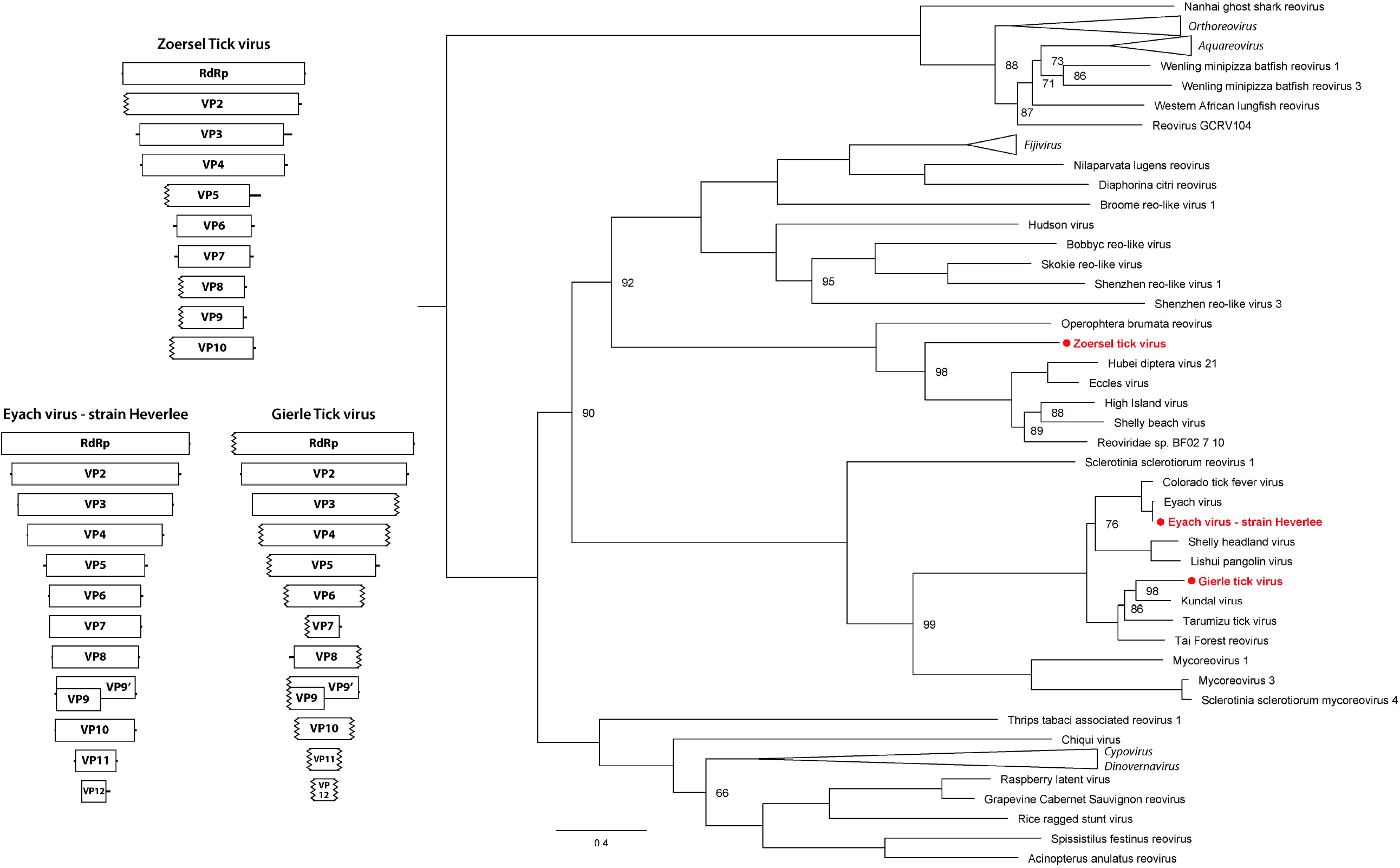
Reovirus phylogeny and genome organization. Left: Genome organization of Eyach virus strain Heverlee, Gierle tick virus and Zoersel tick virus, with segment lengths drawn according to scale. Ribbed lines indicate incomplete genome ends. Right: Maximum-likelihood tree based on the protein sequence of the RdRp of all the (putative) members of the subfamily *Spinareovirinae* for whom such a sequence is available, shows Eyach virus strain Heverlee and Gierle tick virus clustering within the genus *Coltivirus*, with Gierle tick virus representing a novel species. Zoersel tick virus forms a separate branch within the subfamily, alongside several yet unclassified viruses. Branch lengths are drawn according to scale, representing the amount of amino acid substitutions. The numbers at the node indicate the bootstrap support for each node, based on 1000 replicates. Only values <100 are shown. All used accession numbers are provided in Supplementary Table S4.

### Prevalence of selected viruses

To get an indication of the abundance of these novel viruses in the Belgian tick population, we screened all 239 RNA pools by RT-PCR for the presence of the newly discovered viruses mentioned above, as well as the viruses recently discovered in *I. ricinus* from Norway (see [11]) and TBEV. An overview of the screening results is shown in Figure 5 and a table detailing all PCR results is provided in Supplementary Table S1. Except for TBEV, all viruses were detected in one or more pools. Norway luteo-virus 4 was not detected by PCR, but some reads corresponding to this virus were present in one of the Illumina libraries. Presumably, the abundance of this virus was too low and/or the sample quality insufficient to allow detection by PCR. Especially Norway partiti-like virus 1 and Grotenhout virus were highly abundant, being present in >50% of pools screened. In addition, Leuven phlebovirus, Chimay rhabdovirus and Bronnoya virus were also present in a significant portion of samples (17-27% of pools), while all other viruses were only present sporadically (<6%).

**Figure 5:**
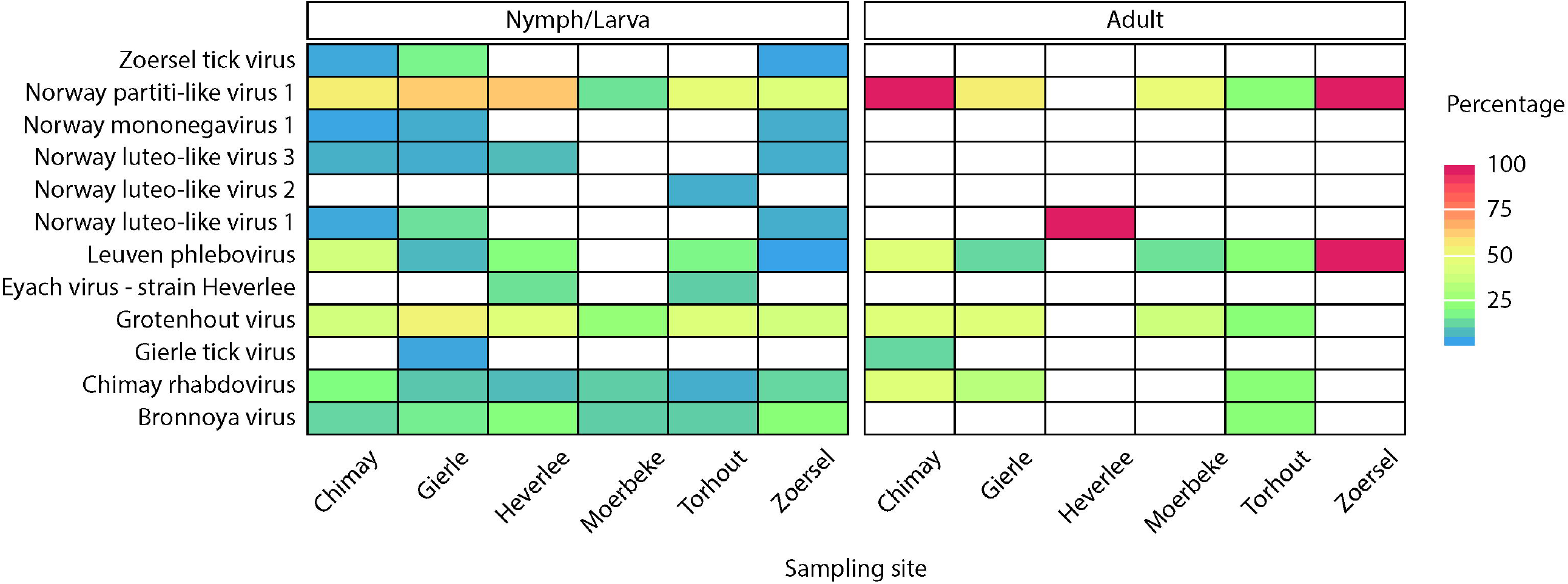
Virus prevalence at the different sampling sites. Overview of all PCR screening results showing no clear correlation of any of the detected viruses with developmental stage/location of origin. TBEV and Norway luteo-like virus 4 were not detected by PCR in any of the screened pools. Percentages shown here are likely over- or under-representations of the true values due to pooling of ticks during extraction and limited counts for some of the developmental stages/locations. A full overview of all screening results can be found in Supplementary Table S1.

## Discussion

Several publications in recent years have provided insight into the diversity of viruses carried by ticks [8-14,21,22]. Although many unknowns remain for most of these viruses in terms of host and geographical range as well as pathological relevance for human, animal and plant health, the discovery of novel viruses is an important first step in expanding our knowledge on viral diversity and viral evolution. As ticks are known vectors of important viral zoonoses, mapping the diversity of viruses carried by ticks can also aid in identifying the origin of yet unidentified or emerging zoonoses. In this study, we report the discovery of several novel viruses found in *I. ricinus*. In addition, we screened >200 tick pools from different sampling locations in Belgium for the presence of these and other, previously described, viruses. While the differences in number of samples per location and the spread of the used sampling sites throughout Belgium do not allow unbiased quantification of the spread of these viruses in the Belgian tick population, the data presented here does provide an indication of virus prevalence. Some viruses are detected in up to 75% of pools, while most are only found sporadically. We detected no TBEV, in line with previous studies screening for TBEV in Belgian ticks [16]. Interestingly, there seems to be no clear correlation between the presence of specific viruses and tick developmental stage or geographical origin, as most viruses are found in both adult and nymph stages, and in several locations. Absence of certain viruses in certain developmental stages/locations can be most likely attributed to under-sampling of those conditions for those specific viruses, although the sample set would need to be expanded significantly to prove this.

Perhaps the most interesting discovery is the finding of two members of the genus *Coltivirus*. This genus currently contains five member species, although putative additional species have been reported [8,33-35]. Two of its members, Colorado tick fever virus and Eyach virus, are known human pathogens capable of causing severe disease [36]. While the other members, Tarumizu tick virus, Tai Forest reovirus and Kundal virus, have not been implicated in human disease, they have been shown to be capable of infecting mammalian and/or human cell lines [37-39]. One of the two coltiviruses reported here appears to be a strain of Eyach virus, sharing >94% nucleotide similarity for all but one segments. To our knowledge, this marks the first detection of Eyach virus, or any known viral human pathogen in the Belgian tick population, further expanding the known geographical range in which this virus is found (see [40]). Interestingly, the fourth segment is markedly dissimilar from known Eyach virus sequences, indicating that some form of reassortment has occurred, hinting at the existence of other, yet to be discovered coltiviruses. The fourth segment encodes for VP4, a protein that contains no known conserved domains and, while conserved among members of the genus *Coltivirus*, seems to share no similarities with any known proteins of other reoviruses. As such, evaluating the importance of this reassortment is difficult, although it is possible that the impact on the virus’ pathogenicity or other characteristics is severe, as even minor sequence variations have been observed to significantly affect coltivirus infections *in vitro* [37]. Unlike this strain of Eyach virus, Gierle tick virus clearly represents a novel species within the genus *Coltivirus*, with all 12 segments sharing <70% nucleotide similarity with other known members of the genus. Although we did not manage to isolate the virus itself, its close phylogenetic relationship to other viruses all known to infect mammalian cells does warrant further research into this virus as a putative human or animal pathogen.

In addition to two coltiviruses, we also identified two reoviruses not belonging to any of the currently recognized genera in the family *Reoviridae*. While we only found traces of one of them, Torhout tick virus appears to be distantly related to Fennes virus, a virus recently identified in ticks found on Antarctic penguins [41]. Conversely, a near-complete genome was reconstructed for Zoersel tick virus. Phylogenetically, this virus is most closely related to a group of viruses for which only six genome segments are known, and, to a lesser extent, the ten-segment containing Operophtera brumata reovirus. Interestingly, several of the ten segments we identified for Zoersel tick virus share (almost) no sequence similarity with any known sequences. Without the minimal sequence similarity between segments 5, 7 and 10, and the corresponding segments of Operophtera brumata reovirus (NC_007559-NC_007568), identifying all ten segments would have been significantly more difficult. Thanks to the availability of this reference, the erroneous conclusion that Zoersel tick virus has only six genome segments, like all viruses with whom it clusters, could be avoided. Interestingly, the nearest neighbours of Zoersel tick virus are the only known reoviruses to have as little as six genome segments. However, the high degree of sequence divergence of the additional segments of Zoersel tick virus raises the concern that these viruses might also have additional segments that are highly divergent. Such divergence could leave certain segments undiscovered by typical BLAST-based approaches, a concern previously raised by some of the groups that discovered these putative six-segmented viruses [42-44]. Re-analyzing these viruses’ sequencing sets with the help of additional related sequences, such as the ones presented here, might help identifying potentially missed segments.

For several of the viruses we detected in our sequencing data, highly similar or related viruses have previously been reported in different tick species from various locations, strengthening the hypothesis that these viruses are truly tick-associated. As noted by Pettersson and colleagues, this association can sometimes be difficult to claim based only on sequencing data, especially if the viruses in question are only found infrequently or in low abundance [11]. Intriguingly, two of the viruses for which they note such low abundance, Norway mononegavirus 1 and Norway luteo-virus 4, are also some of the least-detected viruses in our sample set, although we did find traces of both. Their detection in multiple unrelated datasets might suggest that they are simply low abundant tick-associated viruses, rather than environmental contaminants. In addition to previously identified viruses, we also detected a variety of novel viruses in our data. For some of these viruses, (near-)complete genomes could be reconstructed (see above), albeit most were only present in low abundance, resulting in only small fragments of their genomes being recovered. The low abundance and associated lack of available sequence data make it difficult to confidently classify these viruses as tick-associated, although the similarity of some of them with members of virus families and orders known to contain tick-borne viruses (*Rhabdoviridae, Phenuiviridae, Chuviridae, Orthomyxoviridae* and *Mononegavirales*) do seem to support this association. For many other viruses of which we find traces, which only share similarities with unclassified viruses or viruses belonging to virus lineages with mixed hosts, the origin remains largely unknown. Nonetheless, they do provide further evidence that virus diversity in ticks (and other arthropods) is much higher than until recently assumed. Additional studies are needed to further map this diversity, allowing a better understanding of tick-associated virus evolution. A better characterization of the virus diversity in tick populations can also facilitate studying the spread of known and novel zoonotic viruses, allowing more targeted pre- and post-exposure countermeasures for tick-borne diseases to be taken where needed.

In conclusion, we present the first study looking at virus diversity in *I. ricinus* from Belgium. We show that Belgian ticks carry a wide variety of viruses and that several of these viruses are found in different locations and different tick developmental stages. We also managed to reconstruct (near-)complete genome sequences for several novel viruses, including a new strain of the pathogenic Eyach virus, a new coltivirus species and a highly divergent reovirus that remains to be classified. This research further illustrates that many tick-associated viruses remain to be discovered, and that additional studies are needed to expand our knowledge of virus diversity in ticks and to further characterize these novel viruses.

## Acknowledgments

The authors wish to thank Ana Rita Lopes for excellent technical assistance. BV was supported by a FWO SB grant for strategic basic research of the “Fonds Wetenschappelijk Onderzoek”/Research foundation Flanders [1S28617N].

